# From parental antibiotics to offspring locomotion: L-norvaline links microbiome disruption to intergenerational reprogramming in *Drosophila*

**DOI:** 10.64898/2026.06.17.732813

**Authors:** Jay Dalvi, Ron Schweitzer, Sondra Turjeman, Soliman Khatib, Omry Koren

## Abstract

The maternal microbiome during oogenesis is a critical but underappreciated regulator of offspring physiology. Using *Drosophila melanogaster* as a model, we show that a single, transient antibiotic exposure during oogenesis, without direct offspring exposure, is sufficient to reprogram larval behaviour, gene expression, and metabolism. Adult flies exposed to either ciprofloxacin or vancomycin produced third-instar larvae with significantly reduced crawling speed and a significantly higher prevalence of abnormal locomotion. Transcriptomic profiling revealed widespread downregulation of genes governing energy metabolism, particularly the tricarboxylic acid cycle, as well as cuticle development and protein folding. Metabolomic profiling identified elevation of L-norvaline across both antibiotic groups, and acute parental supplementation with L-norvaline alone recapitulated the observed crawling deficits, directly implicating this metabolite in the phenotype. These findings demonstrate that even brief disruption of the microbiome during oogenesis triggers a cascade of intergenerational physiological changes, establishing this window as a key period of microbiome-dependent offspring programming.

## Introduction

The gut microbiota plays a central role in nutrient metabolism, immune system maturation, and defence against pathogen colonisation^1^. It also influences host behaviour through the gut-brain axis^2–5^. This bidirectional communication is mediated in part by microbial metabolites acting through the immune, endocrine, and nervous systems to influence cognitive function and social interactions^6^. Antibiotics, while indispensable for combating bacterial infections, exert broad, indiscriminate pressure on the gut microbiome, disrupting its equilibrium and potentially precipitating dysbiosis^7^. Such antibiotic-induced dysbiosis has been implicated in adverse outcomes, including compromised maternal immune function and suboptimal offspring immune programming^8^. Antibiotic use during reproduction also has repercussions for both maternal physiology and offspring development through modulation of the maternal gut microbiome and metabolome^9^. *Drosophila* harbour relatively simple but biologically important microbial communities that modulate oogenesis, immunity, metabolism, and lifespan; disrupting these communities with antibiotics therefore has the potential to alter maternal provisioning and indirectly shape offspring phenotypes. Indeed, adult antibiotic exposure in *Drosophila* has been shown to impair oogenesis and reduce fecundity^10,11^, and parental antibiotic exposure can remodel the F1 adult microbiome^12^ with consequences extending into the next generation^13–15^. Yet it is still unclear how acute perturbations during the specific window of oogenesis produce lasting intergenerational effects on offspring physiology. Microbiota alterations have also been linked to changes in behaviour across humans, rodents, and flies, including locomotion, stress responses, learning, and social behaviour^16–22^, but the underlying mechanisms by which microbiome disruption translates into behavioural change remain poorly understood.

*Drosophila melanogaster* has long served as a powerful model organism in molecular genetics, developmental biology, and physiology due to its short life cycle, well-annotated genome, and conserved signalling pathways that parallel those of higher eukaryotes^23^. The reproductive biology of *Drosophila*, particularly oogenesis, involves a tightly orchestrated interplay of genetic, epigenetic, and hormonal factors. Microbial metabolites play an integral part in this process, modulating the activity of juvenile hormone and 20-hydroxyecdysone, both of which are critical for the proliferation of germline stem cells and timely egg maturation^24^. Disruption of the microbiota during oogenesis has also been shown to alter systemic hormonal profiles and accelerate the maternal-to-zygotic transition (MZT) in early embryos^13^. The MZT is a critical phase in which control of developmental processes shifts from maternally deposited transcripts to zygotically transcribed RNA, and it is highly sensitive to the temporal regulation of gene expression and cellular signalling^13^. Consistent with this, recent research has shown that environmental perturbations affect not only the reproductive output of the F0 female but also the developmental trajectory of F1 offspring^13,25–28^. Direct alteration of microbial communities in *Drosophila* has been linked to modifications in feeding behaviour, locomotion, mating preferences, and social interactions^20,29^. Although behavioural assays following parental antibiotic exposure are limited, one study reported that maternal, paternal, and joint parental tetracycline exposure resulted in developmental delays in *Drosophila suzukii* ^30^. A mechanistic understanding of these phenotypes is still lacking.

Understanding how parental antibiotic exposure translates into offspring phenotypes requires integrating information across biological scales. Transcriptomic profiling has revealed how early-life microbiome disruption alters gene regulatory networks governing immunity, metabolism, and development in *Drosophila* offspring^13,28,31^. Metabolomic approaches have further shown that the maternal microbiome shapes the pool of maternally provisioned metabolites in the egg, including amino acids, lipids, and tricarboxylic acid (TCA) cycle intermediates that fuel early embryogenesis and larval physiology^32–34^. Behaviour emerges as the functional readout of these molecular disruptions: locomotion in *Drosophila* larvae is sensitive to perturbations in energy metabolism, neuromuscular signaling, and cuticle integrity^35–38^. Yet how antibiotic exposure during the oogenesis window cascades through these biological layers, from disrupted maternal provisioning to altered gene expression and metabolic reprogramming, to aberrant larval physiology, remains unresolved. Clarifying this chain of causation in a tractable model organism is essential for understanding whether the intergenerational consequences of parental antibiotic exposure reflect a generalised stress response or a targeted disruption of microbiome-dependent developmental programmes.

Here, we exposed *D. melanogaster* parental flies to either ciprofloxacin or vancomycin during oogenesis and profiled the F1 larvae through behavioural assays, RNA sequencing, and metabolomic profiling (Fig. 1a). To test the mechanistic contribution of a key metabolite identified in our analysis, we also performed a targeted L-norvaline supplementation experiment using the same experimental design. Ciprofloxacin is a broad-spectrum fluoroquinolone antibiotic that inhibits bacterial DNA gyrase and topoisomerase IV, preventing DNA replication, and is effective against many Gram-negative and some Gram-positive bacteria^39^. Vancomycin is a glycopeptide antibiotic that inhibits cell wall synthesis in Gram-positive bacteria by binding to D-Ala-D-Ala termini of peptidoglycan precursors^40^. Through our multi-omics approach, we revealed alterations in larval locomotion, gene expression, and metabolism in offspring of antibiotic-exposed flies, with elevated L-norvaline emerging as a key contributing metabolite.

**Figure 1.**
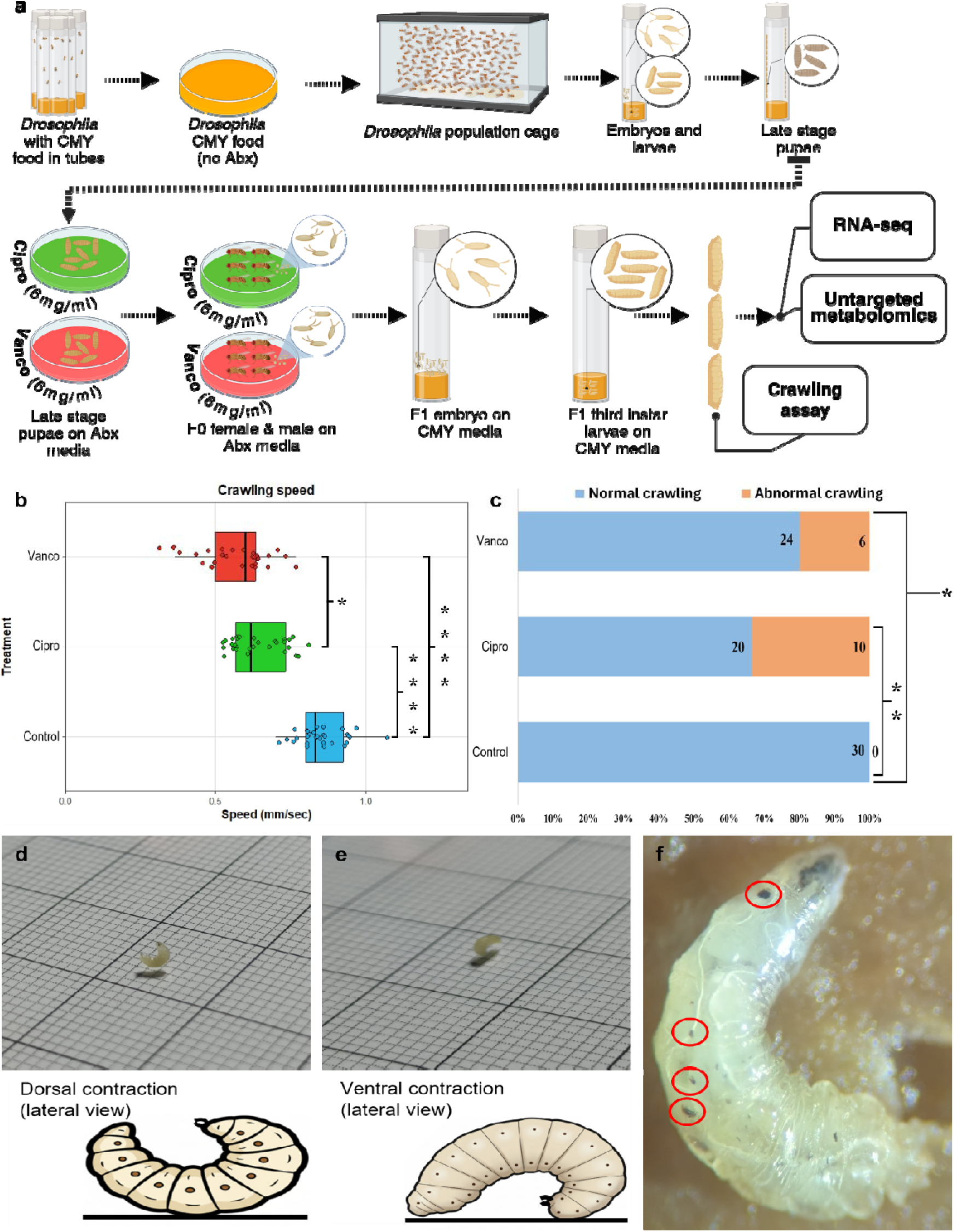
Experimental design and effects of parental antibiotic exposure on larval phenotype. (a) Schematic overview of the experimental workflow. CMY: cornmeal, molasses, yeast; Abx: antibiotics. (b) Crawling speed of F1 third-instar larvae from control, ciprofloxacin (Cipro), or vancomycin (Vanco) treated parents. Each point is one larva; boxplots show median and interquartile range; Asterisks denote significance versus control (data in Table S1). (c) Percentage of larvae exhibiting normal (blue) or abnormal (orange) crawling per treatment group; numbers within bars indicate sample sizes and asterisks indicate significant differences from control. (d,e) Representative examples of larvae with abnormal dorsal (d) or ventral (e) contractions during locomotion, with corresponding schematics. (f) Magnified image of a third-instar larva of ciprofloxacin-treated parents showing melanotic masses (circled in red). * = q < 0.05; ** = q < 0.01; **** = q < 0.0001.

## Results

### Parental antibiotic exposure impairs offspring locomotor behaviour

Following parental antibiotic exposure for 24-30 h spanning the oogenesis window, we assessed the locomotor behaviour of F1 third-instar larvae. Quantitative analysis of crawling speed revealed significant differences between treatment groups. Control larvae crawled significantly faster (0.852 ± 0.079 mm/s, mean±SD) than larvae from ciprofloxacin-treated parents (0.639 ± 0.088 mm/s) and from vancomycin-treated parents (0.563 ± 0.116 mm/s; both q < 0.0001 versus control; Fig. 1b). Among the two antibiotic-treated groups, larvae of vancomycin-treated parents crawled significantly more slowly than those from ciprofloxacin-treated parents (q = 0.027; Fig. 1b).

We also systematically monitored larvae for morphological and behavioural abnormalities during the crawling assay. We observed abnormal crawling phenotypes in 33% of larvae of ciprofloxacin-exposed parents (q = 0.0016) and 20% of larvae of vancomycin-exposed parents (q = 0.0237), while no such abnormalities were detected in control larvae (Fig. 1c). These behavioural abnormalities were categorised based on the pattern of body contractions, with normal larvae displaying coordinated peristaltic waves and abnormal larvae showing either predominantly dorsal (Fig. 1d) or ventral contractions (Fig. 1e). Melanotic masses were also observed in offspring from both antibiotic treatment groups at the third-instar larval stage but not in control larvae; the difference in prevalence did not reach statistical significance (Fig. 1f).

### Transcriptomic analysis reveals widespread and treatment-specific changes in gene expression

To elucidate the molecular mechanisms underlying the observed behavioural phenotypes, we performed bulk RNA sequencing (RNA-seq) on third-instar larvae from all treatment groups. Principal component analysis (PCA) of the normalised gene expression data demonstrated clear treatment-specific separation. When comparing pools of larvae of ciprofloxacin-exposed parents to pools of control larvae, PC1 explained 35.46% and PC2 explained 21.74% of the total variance (Fig. 2a); in the vancomycin-control comparison, PC1 and PC2 explained 41.64% and 16.52%, respectively (Fig. 2b).

**Figure 2:**
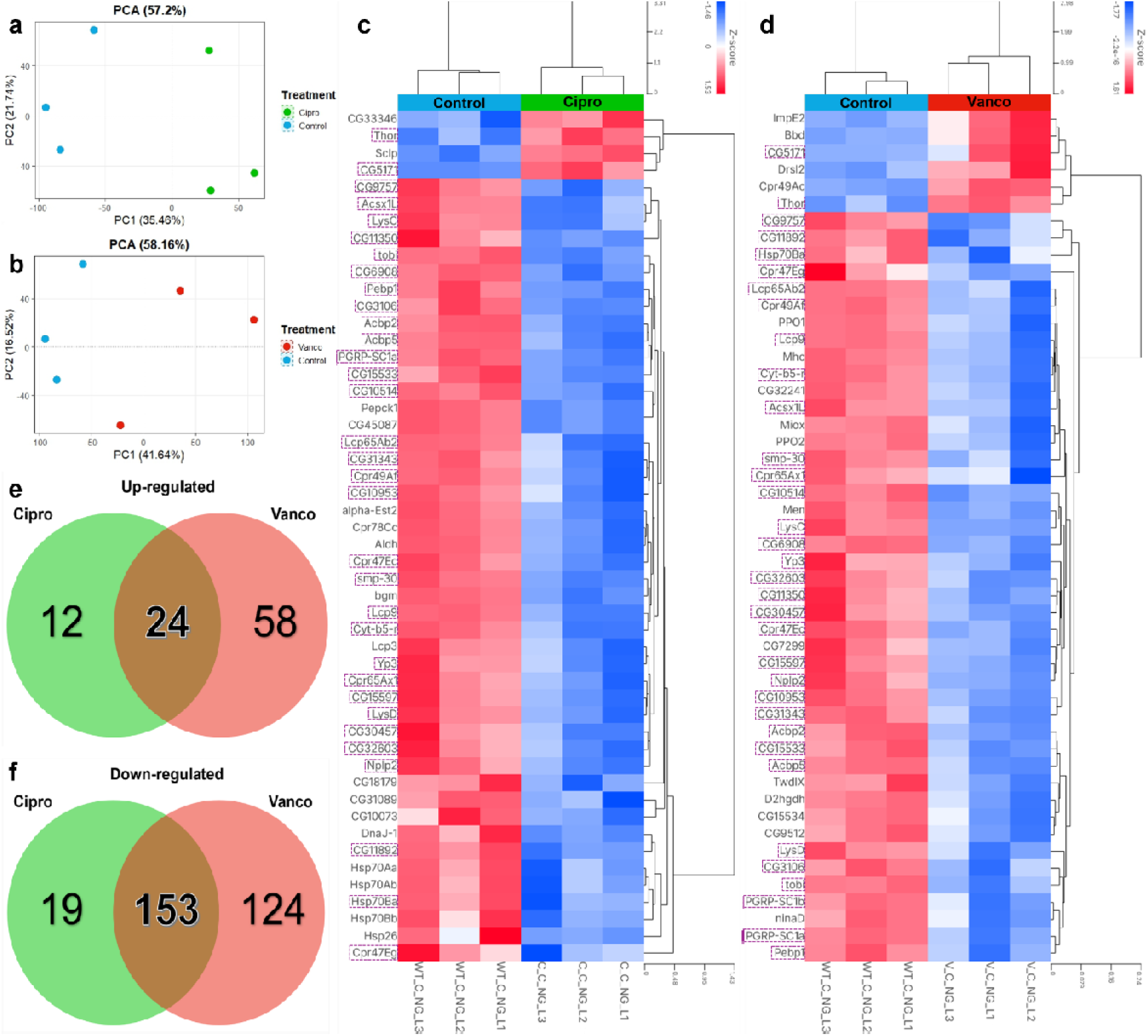
**Transcriptional response in F1 larvae following parental antibiotic exposure during oogenesis**. (a,b) PCA of RNA-seq profiles from pools of 20 F1 third-instar larvae (n = 3 pools per treatment group), showing treatment-specific clustering of larvae of (a) ciprofloxacin-exposed and (b) vancomycin-exposed parents relative to controls. (c,d) Heatmaps of the top 50 DEGs in the (c) ciprofloxacin and (d) vancomycin groups versus the control, with red indicating relative upregulation and blue indicating relative downregulation (z-scores of log-normalised expression values) across individual samples and purple dashed boxes highlighting shared DEGs between the treatment groups. (e,f) Venn diagrams illustrating the overlap of significantly (e) upregulated and (f) downregulated genes between the ciprofloxacin and vancomycin treatment groups, highlighting shared and treatment-specific DEGs.

Differential gene expression analysis, with Benjamini-Hochberg false discovery rate (FDR) corrections and an absolute fold change (FC) of ≥2, identified significant transcriptional changes in both treatment groups. In larvae of ciprofloxacin-exposed parents, we identified 208 significantly differentially expressed genes (DEGs) relative to controls, comprising 36 upregulated and 172 downregulated genes (Fig. 2c). Larvae of vancomycin-exposed parents exhibited a more pronounced transcriptional response, with 359 DEGs, including 82 upregulated and 277 downregulated genes (Fig. 2d). Comparative analysis revealed 24 genes commonly upregulated (Fig. 2e) and 153 genes commonly downregulated (Fig. 2f) across both antibiotic treatment groups.

### Gene ontology and KEGG pathway analysis reveal disrupted biological processes critical for development and metabolism

To gain functional insights into the observed transcriptional changes, we performed Gene Ontology (GO) enrichment analysis on DEGs using ShinyGO, iDEP and g:Profiler^41–43^. This analysis revealed distinct patterns of pathway disruption between the two antibiotic treatment groups across several key annotation classes, including GO biological processes (GO-BP), cellular components (GO-CC) and molecular functions (GO-MF), as well as human ortholog pathways (HP) (Fig. S1).

Focusing on GO-BP terms, the downregulated genes in larvae of ciprofloxacin-treated parents were enriched for developmental and metabolic processes including cuticle development, energy production, protein folding and response to environmental stress (Fig. 3a). In larvae of vancomycin-treated parents, downregulated genes were associated with cellular structure and development, immune responses, energy production and protein folding (Fig. 3b). KEGG pathway analysis revealed that multiple metabolic pathways were downregulated in both treatment groups, particularly pathways involved in energy production (Fig. 3c,d). Key TCA cycle enzymes, including citrate synthase (*kdn*) and malate dehydrogenase 2 (*Mdh2*), were downregulated in both treatment groups (ciprofloxacin: *kdn* q = 0.0019, *Mdh2* q = 0.0021; vancomycin: *kdn* q = 0.0009, *Mdh2* q = 0.0009, Fig. 3e,f). Phosphoenolpyruvate carboxykinase 1 (*Pepck1*), a gluconeogenic enzyme that operates at the interface of the TCA cycle and gluconeogenesis, was also downregulated in both treatments, though the FC did not exceed the threshold of >2 for the vancomycin treatment group (ciprofloxacin q < 0.0001, FC: −2.645; vancomycin: q = 0.0098, FC: −1.910; Fig. 3g). In addition, expression of aldehyde dehydrogenase (*Aldh*) was significantly downregulated in both ciprofloxacin (q < 0.0001) and vancomycin (q < 0.0001) treatment groups (Table S2, S3).

**Figure 3:**
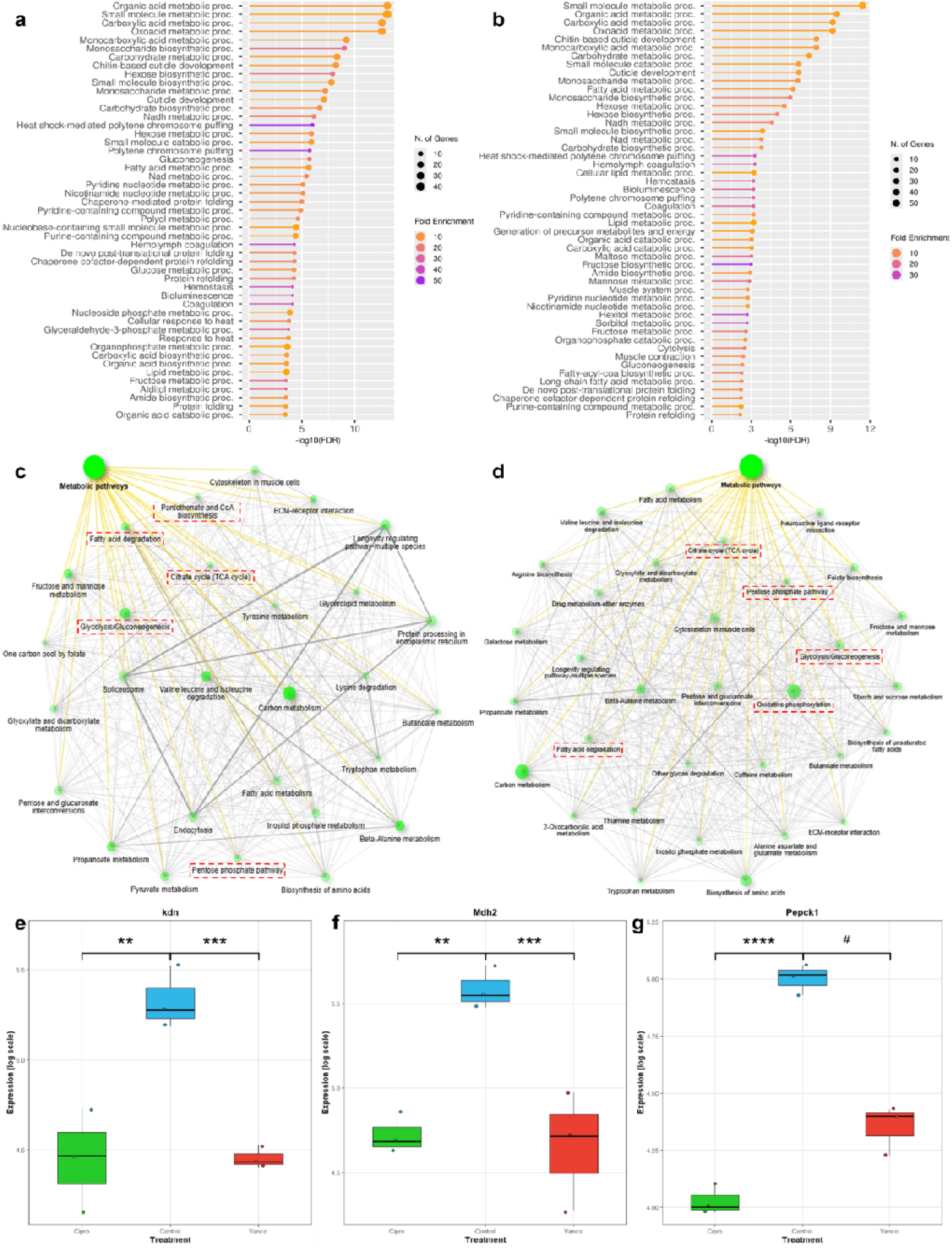
Treatment-specific pathway enrichment and metabolic gene regulation in F1 larvae. (a,b) Top enriched GO biological process (GO-BP) terms among downregulated DEGs in larvae of (a) ciprofloxacin-exposed and (b) vancomycin-exposed parents versus controls, ranked by enrichment significance. The bar length represents the −log10 FDR values on the x-axis; the colour represents the fold enrichment, and the circle size represents the number of genes in the pathway. (c,d) Network visualisation of enriched KEGG pathways among downregulated DEGs. Each node represents an individual pathway, with edges indicating shared gene membership between pathways for larvae of (c) ciprofloxacin and (d) vancomycin-exposed parents versus controls. Each green dot represents an individual pathway, and lines indicate interconnectivity of the pathways; metabolic modules including the TCA cycle, glycolysis/gluconeogenesis, pentose phosphate, fatty acid-related pathways and oxidative phosphorylation are highlighted with red boxes (modules vary by panel). (e-g) Boxplots of log-scaled expression of (e) kdn, (f) Mdh2, and (g) Pepck1 in ciprofloxacin, control and vancomycin groups, with q-values from DESeq2 analysis with Benjamini–Hochberg FDR correction (n= 3 pools per treatment group). Note that the fold change of Pepck1 in the control vs vancomycin comparison did not pass the threshold of 2 and is therefore denoted with # despite a significant q-value. ** = q < 0.01; *** = q < 0.001; **** = q < 0.0001.

Among upregulated genes, enriched GO-BP terms in both treatment groups included the catecholamine metabolic process, cuticle pigmentation, amine biosynthesis, and amino acid metabolism (Fig. S2). The extracellular matrix receptor interaction pathway was downregulated in both treatment groups, including the collagen-encoding genes *vkg* (ciprofloxacin: q = 0.0090; vancomycin: q = 0.0005) and *Col4a1* (ciprofloxacin: q = 0.0054; vancomycin: q = 0.0005; Tables S2, S3). Downregulation was also evident across KEGG metabolic pathways, GO cellular component terms (notably extracellular matrix and muscle-related components), and GO molecular function terms, as detailed in Fig. S3–S5.

### Metabolomic profiling reveals altered metabolite abundance and disrupted biochemical pathways

To complement our transcriptomic findings, we performed untargeted HPLC-MS/MS metabolomic profiling on pools of larvae from all treatment groups. PCA of metabolomic profiles revealed clear separation between larvae of the two antibiotic-treated groups and controls. In the ciprofloxacin versus control comparison, PC1 and PC2 explained 81.5% and 11.7% of the total variance, respectively (Fig. 4a). In the vancomycin versus control comparison, PC1 and PC2 explained 89.8% and 6.1% of the variance, respectively (Fig. 4b).

**Figure 4:**
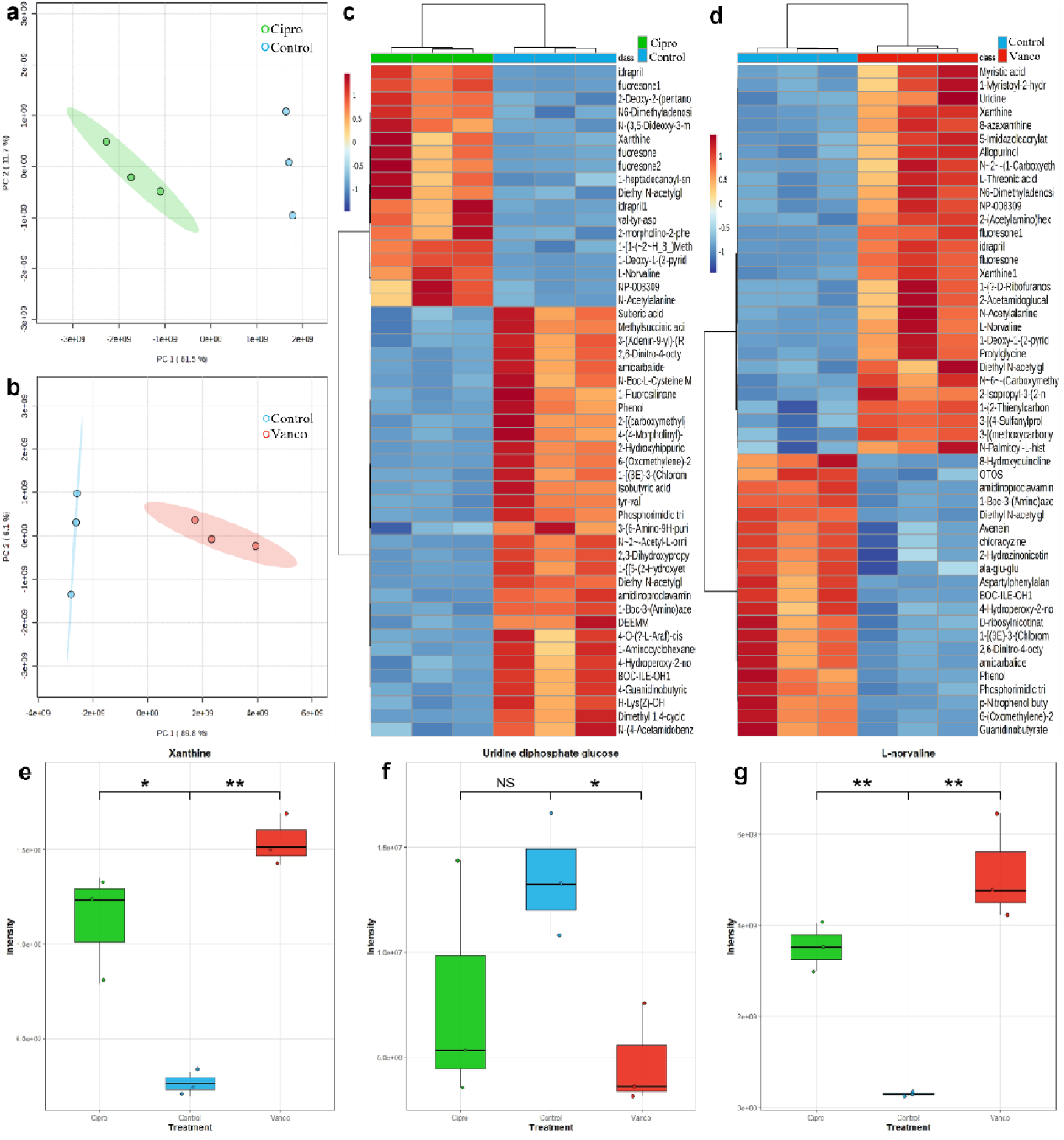
Untargeted metabolomic profiling of F1 larvae following parental antibiotic exposure. (a,b) PCA of metabolite profiles from ciprofloxacin (a) and vancomycin (b) groups versus controls (n = 3 per group). (c,d) Heatmaps of the top 50 significantly altered metabolites ranked by FDR-corrected p-value in ciprofloxacin (c) and vancomycin (d) versus control. Each column is one biological pool (n = 3 pools per treatment group); colour scale represents normalised peak area intensity (z-score) (e-g) Boxplots of (e) xanthine, (f) uridine diphosphate glucose, and (g) L-norvaline levels were significantly different in one or both treatment groups compared to the control group (n = 3 per group). NS = not significant; * = q < 0.05; ** = q < 0.01.

In the larvae of ciprofloxacin-treated parents, we identified 70 significantly altered metabolites compared to controls, comprising 32 elevated and 38 reduced compounds (Table S4). Larvae of vancomycin-treated parents showed significant metabolic disruption, with 67 significantly altered metabolites comprising 36 elevated and 31 reduced compounds (Table S5). Hierarchical clustering heatmaps of the top 50 most significantly altered metabolites revealed clear treatment-specific metabolic signatures (Fig. 4c,d).

Xanthine was significantly elevated in both treatment groups (ciprofloxacin: q = 0.02; vancomycin: q = 0.002) (Fig. 4e). Conversely, uridine diphosphate glucose levels were significantly decreased in larvae of vancomycin-treated parents (q = 0.034), while no significant change was detected in larvae of ciprofloxacin-exposed parents (Fig. 4f). Notably, L-norvaline was significantly elevated in larvae from both ciprofloxacin- and vancomycin-exposed parents (ciprofloxacin: q = 0.0049; vancomycin: q = 0.0093) (Fig. 4g).

### Parental L-norvaline supplementation phenocopies the locomotor deficits induced by antibiotic exposure

Based on our finding that L-norvaline was elevated in larvae of both antibiotic-exposed groups, and given prior evidence linking this metabolite to locomotor function^44^, we supplemented F0 parental diets with L-norvaline during oogenesis to directly test its contribution to the observed F1 crawling deficits. Parental L-norvaline supplementation (1 mg/ml) following the same experimental design as the antibiotic exposures (Fig. 1a) resulted in a significant reduction in larval crawling speed (0.517 ± 0.111 mm/s) relative to the matched control group (Control-2: 0.850 ± 0.057 mm/s; q < 0.0001). This speed was comparable to that observed in larvae of vancomycin-exposed parents (q = 0.173) and significantly lower than that of ciprofloxacin-exposed larvae (q = 0.0012; Fig. 5). Abnormal crawling phenotypes were also observed in larvae of L-norvaline-supplemented parents, though this did not reach statistical significance (Fisher’s exact test, q = 1).

**Figure 5:**
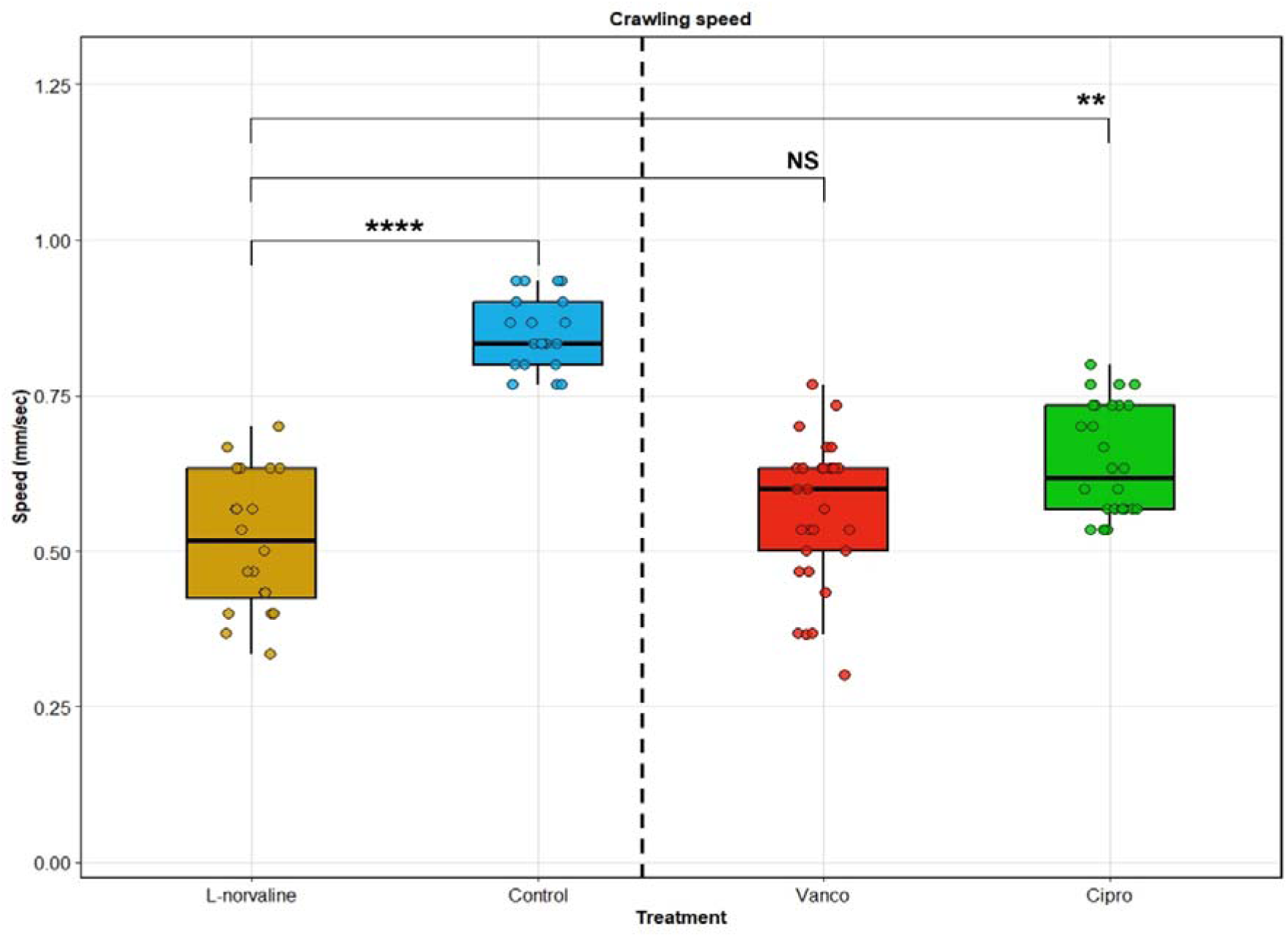
F0 L-norvaline supplementation phenocopies the F1 crawling deficits observed in offspring of antibiotic-exposed parents. Boxplots showing crawling speed in larvae from L-norvaline-supplemented parents and the corresponding control group (n = 20 larvae per group), as well as larvae of vancomycin- and ciprofloxacin-treated parents (n = 30 larvae per group), with individual data points overlaid. NS= not significant; **= q < 0.01; **** = q < 0.0001, two-tailed Kruskal-Wallis test. Note that the L-norvaline and corresponding Control groups (left of dashed line) were part of a separate experiment conducted under an identical design to the antibiotic-exposure experiment (right of dashed line, see Fig. 1).

## Discussion

Transient parental antibiotic exposure spanning the *Drosophila* oogenesis window is sufficient to reprogram offspring physiology across behavioural, transcriptomic, and metabolomic levels without any direct antibiotic exposure of the larvae themselves. Acute parental exposure to both ciprofloxacin and vancomycin produced convergent disruptions in larval locomotion, energy metabolism, and cuticle development, demonstrating that the F0 microbiome during oogenesis is critically important for normal offspring physiology, and that its disruption has consequences extending beyond the exposure window.

One of the clearest outcomes was the abnormal crawling behaviour in larvae of parents exposed to antibiotics. Previous research has described similar effects following direct antibiotic exposure in *Drosophila* and other species, including rodents and a nonhuman primate, ranging from teratogenicity and developmental delays to altered lifespan, social interactions and anxiety-like behaviour^15,27,45–47^.

While F0 exposure to both antibiotics induced intergenerational effects on offspring neuromuscular function, vancomycin had a significantly more pronounced effect on larval crawling speed than ciprofloxacin (Table S1). This may reflect the different mechanisms of action of these antibiotics or their differential effects on parental physiology during oogenesis. The shared abnormal crawling phenotype likely reflects broader systemic disruptions in the neural circuits controlling coordinated muscle contraction or defects in neuromuscular junction function^36,37^.

RNA sequencing of third-instar larvae revealed widespread transcriptomic dysregulation in both antibiotic treatment groups compared to the control, with downregulated genes substantially outnumbering upregulated genes. GO-BP enrichment analysis revealed that both antibiotic treatments resulted in downregulation of cuticle development, energy production, protein folding and environmental stress responses. In particular, key TCA cycle enzymes and associated metabolic pathways were downregulated in both groups. This metabolic reprogramming likely contributes to the observed locomotor deficits and morphological abnormalities, as developing larvae require substantial energy resources for growth and moulting^48^. Downregulation of *vkg* and *Col4a1*, which encode type IV collagen subunits essential for extracellular matrix integrity and basement membrane assembly, was also observed in both treatment groups. Compromised cuticle integrity and basement membrane structure could directly contribute to the observed locomotor deficits by impairing the mechanical properties required for coordinated peristaltic movement. Disrupted epithelial architecture and immune activation may underlie the morphological abnormalities, including melanotic masses and aberrant body contractions, consistent with parental antibiotic exposure predisposing offspring to inflammatory responses or tissue dysfunction^49,50^. The *Drosophila Aldh* gene, which was significantly downregulated in both treatment groups, has been reported to be suppressed under conditions of microbial dysbiosis, with inhibitory effects on oogenesis^13^.

While gene suppression was widespread, both antibiotics also induced upregulation of distinct metabolic pathways. F0 ciprofloxacin exposure upregulated genes associated with purine biosynthesis, suggesting a potential compensatory response. This upregulation may reflect cellular efforts to sustain DNA repair and replication processes under ciprofloxacin-induced stress, a phenomenon that has also been observed in human lymphocytes^51^. In the ciprofloxacin group, genes associated with glycolysis, the pentose phosphate pathway, and the TCA cycle were downregulated, whereas genes involved in *de novo* purine synthesis were upregulated. These pathways are functionally interconnected through shared metabolic intermediates, which may indicate a redistribution of metabolic flux toward *de novo* purine biosynthesis^52^ and increased endogenous nucleic acid turnover, leading to enhanced purine degradation^53,54^. In contrast, the vancomycin treatment was associated with enhanced expression of genes involved in catecholamine and amino acid biosynthesis. The upregulation of catecholamine metabolism is especially intriguing, as catecholamines play a crucial role in *Drosophila* behaviour including locomotion, learning, and courtship^55–57^, and this upregulation may represent compensatory mechanisms for the observed locomotor impairments. Both antibiotic treatments showed convergent downregulation of energy production and protein folding pathways, suggesting that these represent core mechanisms of antibiotic-induced intergenerational physiological disruption. Similar intergenerational effects have been reported following maternal antibiotic exposure in other species, with offspring showing altered growth and survival^58^, further supporting the view that antibiotic-induced dysbiosis during reproduction has broad consequences for the next generation.

The metabolic alterations observed in both groups demonstrate that the effects of parental antibiotic exposure extend beyond transcriptional reprogramming to influence fundamental biochemical processes. The metabolomic profiles of larvae from antibiotic-exposed parents were markedly distinct from controls, reflecting broad disruption of amino acid, purine, and nucleotide metabolism. Both treatment groups exhibited elevated levels of xanthine and L-norvaline, indicating that these metabolic perturbations represent common responses to parental antibiotic exposure regardless of the specific antibiotic used. Xanthine, a purine degradation product that accumulates during cellular stress and energy depletion^59–61^, was elevated in larvae of both antibiotic-exposed groups, consistent with increased nucleotide turnover and metabolic stress. Elevated L-norvaline, a non-proteinogenic amino acid that inhibits arginase and increases nitric oxide production, may indicate disrupted amino acid synthesis pathways or cellular stress responses^62^. Consistent with these findings and follow-up F0 supplementation experiment, L-norvaline supplementation has been reported to reduce locomotion in zebrafish larvae^44^, further supporting its role in larval locomotion. Uridine diphosphate glucose, the primary sugar donor for UDP-glycosyltransferase enzymes that drive detoxification and diverse physiological processes in *Drosophila*^63^, was significantly reduced in vancomycin-treated offspring, suggesting impaired conjugation capacity and potentially compromised xenobiotic clearance. Collectively, these findings reinforce the view that antibiotics administered during the sensitive window of oogenesis do more than simply clear bacteria; they remove symbiotic signals necessary for normal egg development. The ensuing disruption of maternally provisioned metabolic and molecular signals likely predisposes offspring to physiological alterations that emerge during embryogenesis and persist throughout larval development.

Several limitations of this study merit consideration. First, both male and female parental flies were housed on antibiotic-supplemented food, meaning we cannot exclude a contribution of paternal microbiome disruption to the observed offspring phenotypes. Future experiments using sex-segregated exposure designs would help disentangle maternal and paternal contributions. Second, offspring were characterized at a single developmental stage; whether these physiological alterations persist into later larval instars or adulthood remains to be determined. Similarly, future research should also examine the multigenerational consequences of such acute exposures, through the F2 or F3 generations.

Our findings align with growing evidence that the maternal microbiota plays an essential role in shaping embryonic development, immune programming, and behaviour through microbe-host metabolic signalling^64,65^. While *D. melanogaster* does not have placental architecture, maternal provisioning of microbial metabolites and epigenetic cues during oogenesis and embryogenesis is critical for proper development^24,66^. Antibiotic-induced dysbiosis in the F0 generation during this window likely interferes with microbial contributions to nutrient sensing, hormonal signalling (e.g., ecdysone, juvenile hormone) and transcriptional regulation^67^. Furthermore, antibiotic use during pregnancy has been associated with neurodevelopmental disorders, altered immune function, and behavioural changes in offspring, as shown in rodent and human models^9,18,68–72^. This study adds to the growing body of evidence demonstrating that even indirect antibiotic exposure, limited to parental intake, can elicit broad and systemic changes in the physiology of the next generation.

## Methods

### Fly stocks and culture

Fly stocks (Oregon-R) were obtained from Bloomington *Drosophila* Stock Center (Bloomington, IN, USA). The flies were maintained in 50 ml Bio-Reaction tubes (RED BRT010051-ZX, Alex Red, Israel) containing approximately 7 ml of cornmeal, molasses, yeast (CMY) medium. The recipe was as follows: 6.5 g agar, 76 g cornmeal, 76 g molasses, 50 g dry yeast and 1 l distilled water were mixed and boiled for 30 min with constant stirring and then autoclaved. After autoclaving and cooling, 1.4 g nipagin (H5501, Sigma-Aldrich, St. Louis, MO) dissolved in 7.6 ml 100% ethanol, and 4 ml 99% propionic acid were added separately. The flies were raised under a 12 h light:12 h dark cycle at 25 ^°^C and 60-70% relative humidity in an incubator unless stated otherwise. Once the initial colony reached approximately 300 flies, they were transferred to a population cage made of cast acrylic (61×30×41 cm). The flies in the population cage were maintained on CMY medium in 10 cm petri dishes (Fig. 1a). Petri dishes were replaced every two days; eggs and larvae were harvested from the used dishes, and the adult flies were transferred to the 50 ml Bio-Reaction tubes with CMY medium. The cage population was maintained by transferring new flies twice a week into the cage.

### Experimental design

A fresh 10 cm petri dish with CMY medium was kept in the *Drosophila* population cage for 1 h to collect a synchronised cohort of eggs. The eggs were transferred to 50 ml Bio-Reaction tubes with CMY medium. When the larvae reached the late pupal stage, they were transferred to 10 cm petri dishes with CMY medium supplemented with either ciprofloxacin (6 mg/ml; 17850-25G-F, Sigma-Aldrich, St. Louis, MO) or vancomycin (6mg/ml; V-200-25, GoldBio, St. Louis, MO, USA)^73^. In a separate experiment conducted under the same design, L-norvaline (1 mg/ml; N7627, Sigma-Aldrich, St. Louis, MO) was used in place of antibiotics. In each experiment, control larvae were transferred to and raised on dishes with only CMY medium. Both male and female pupae were placed on antibiotic-containing dishes, allowed to eclose under the same conditions, and maintained on the same petri dishes for 24-30 h. The eggs laid by the F0 generation flies were collected and carefully rinsed with sterile phosphate-buffered saline. These eggs were then transferred to 50 ml tubes containing CMY medium. After hatching and maturation to the third-instar larval stage, the larval crawling behavioural test was performed^74^.

### Locomotion assay

The larval crawling assay was used to assess locomotor function and neuromuscular coordination. The assay was performed as previously described^75^ with the following modifications. Larval crawling assays were performed on 2% water agar in 10 cm dishes placed over graph paper (1 mm grid) for distance measurement. Individual third-instar larvae were recorded crawling for 10 s without interruption in three separate trials (technical triplicates), using a Panasonic HC-V770 video camera. Distance was measured by the number of gridlines crossed. Crawling speed (mm/s) was calculated for 30 larvae per group in the antibiotic exposure experiment and 20 larvae per group in the L-norvaline experiment. The sample size differed in the L-norvaline experiment because not all larvae crawled continuously for 10 s, which was the criterion for inclusion in the crawling-speed analysis. In addition to the behaviourally phenotyped larvae, additional larvae of treatment-exposed parents were collected and snap-frozen in liquid nitrogen and stored at −80°C until transcriptomic and metabolomic processing. The average speed per larva was calculated as the mean of triplicate measurements per individual, and statistical significance between groups was assessed using a Kruskal–Wallis test; results are reported as mean ± standard deviation.

### Observation of altered crawling phenotypes

Larvae were systematically observed for morphological and behavioural deviations from the control phenotype during the crawling assay, and all abnormalities were recorded. This enabled documentation of both gross abnormalities and subtle locomotor changes not captured by molecular analyses alone. Fisher’s exact tests (pairwise) were used to compare each antibiotic-exposed group to the control group and to compare the L-norvaline-exposed group to its matched control.

### Larval transcriptomics

#### RNA extraction and sequencing

RNA was extracted from pools of 20 snap-frozen whole third-instar larvae (three pools per treatment group) using TRI reagent (T9424, Sigma-Aldrich, St. Louis, MO) and 1-Bromo-3-chloropropane (BCP) (B9673, Sigma-Aldrich, St. Louis, MO) following the protocol outlined previously^31,76^ with modifications. The modification consisted of adding β-mercaptoethanol (10 μl) (444203, Sigma-Aldrich, St. Louis, MO) along with TRI reagent (500 μl) and BCP (100 μl). After vortexing and centrifugation (14,000 rpm, 15 min, 4°C), the aqueous phase was transferred to fresh tubes. RNA was precipitated by adding 200 μl pre-chilled isopropanol, vortexing, and incubating at −20 °C for 5 h. The RNA samples were then shipped to Procomcure Biotech NGS, Austria for mRNA sequencing at a sequencing depth of 15 million paired-end reads on an Illumina NovaSeq platform in a 2x150 bp configuration. RNA-seq libraries were prepared using poly-A selection.

#### Data QC and alignment

Unaligned adapter-trimmed FASTQ files were uploaded to the PARTEK server for analysis^77^. Quality control of the reads was performed prior to alignment, and the STAR aligner^78^ tool was used to align the obtained sequences with the FlyBase *Drosophila* reference genome (r6.66)^79^. Post-alignment QC confirmed high mapping rates across all samples (mean alignment rate: 87.55%).

#### Transcriptome analysis

Following quality control and preprocessing, read counts were quantified using the ‘Quantify to annotation model (Partek E/M)’, yielding expression estimates for 17,597 genes across all samples. Genes with a maximum count of ≤ 30.0 across all samples were removed, and the remaining counts were normalised in three steps: i) N + 0.1 (where N = number of counts), ii) Trimmed Mean of M-values, and iii) Log(e). Differential expression analysis was conducted with DESeq2^80^, comparing ciprofloxacin-treated versus control larvae and vancomycin-treated versus control larvae. To correct for multiple comparisons, a false discovery rate (FDR)-correction was applied, and genes with a corrected q-value of ≤ 0.05 and an absolute fold change ≥2 were considered significant. ShinyGO^41^ was used for individual GO & KEGG pathway visualization; iDEP^42^ was used for KEGG pathway analysis; g:Profiler^43^ was used to visualize overall dysregulation across multiple GO and KEGG pathways together from the gene enrichment analysis data^41–43^.

### Larval metabolomics

#### Total metabolite extraction

Metabolites were extracted from whole third-instar larvae following the protocol outlined previously^31,76^ with modifications as follows. Twenty larvae per pool (three pools per treatment) were homogenized in bead tubes (A29158, Thermo Fisher, Waltham, MA) with 600 µl methanol:DDW (80:20) in a Mini Bead Beater (BioSpec Products, Bartlesville, OK) for 2 min. Samples were sonicated on ice in an ultrasonic bath (97043-936, VWR, Radnor, PA, USA) for 10 min, vortexed for 2 min, then centrifuged at 12,000 rpm for 10 min at 4 °C. Five hundred microliters of supernatant were dried in a SpeedVac (Thermo Fisher, Waltham, MA) and resuspended in 55 µl methanol:DDW (50:50). The resuspended extract was then transferred to HPLC inserts (702824, Macherey-Nagel, Düren, Germany) placed inside the microcentrifuge tubes and centrifuged at 4,000 rpm for 10 min at 4 °C. Prior to the final centrifugation step, five microliters from each sample were pooled for quality control. Inserts were then placed into HPLC vials (70213, Macherey-Nagel, Düren, Germany) and stored at – 80 °C until untargeted HPLC-MS/MS metabolomics analysis.

#### HPLC analysis

Extracts (5 μL) were injected into a UHPLC–PDA system (Dionex Ultimate 3000, Thermo Fisher, Waltham, MA) with a ZORBAX Eclipse Plus C18 column (3.0 × 100 mm, 1.8 μm). The mobile phase was DDW + 0.1% formic acid (A) and acetonitrile + 0.1% formic acid (B) at 0.4 mL/min, 30 °C. The gradient was as follows: 2→30% B over 4 min, 30→40% B over 1 min, isocratic at 40% B for 3 min, 40→50% B over 6 min, 50→55% B over 4 min, 55→95% B over 5 min, isocratic at 98% B for 6 min, then return to 5% B over 2 min, and 5 min equilibration.

#### MS/MS analysis

Mass spectrometry analysis was performed with a Heated Electrospray ionisation (HESI-II) source connected to a Q Exactive™ Plus Hybrid Quadrupole-Orbitrap™ Mass Spectrometer (Thermo Fisher, Waltham, MA). HESI capillary voltage was set to 3,900 V in positive mode and 3,500 V in negative mode, capillary temperature to 350 °C, and gas temperature to 350 °C. Sheath gas flow was set to 35 ml/min, auxiliary gas flow to 10 ml/min, and sweep gas flow to 1 ml/min. Mass spectra (m/z 67–1,000) were acquired in negative and positive-ion modes with high resolution (FWHM = 70,000). For MS/MS analysis, normalised collision energy was set to 20, 30 and 50 eV.

#### Metabolome data preprocessing and statistical analysis

Peak determination and peak area integration were performed with Compound Discoverer SP 3.3 (Thermo Fisher Scientific, Version 3.3.0.305). Automatic integration was manually inspected and corrected if necessary. For compound identification and annotation, the MzCloud database^81^ was queried using MS/MS data, and the ChemSpider^82^ and MassList databases were queried using the MS/MS data and HRMS (High-Resolution Mass Spectrometry) data. Remaining compounds were annotated by their molecular formula using the predicted-composition feature in Compound Discoverer SP 3.3 or as m/z. A score for each possible structure was calculated using the mzLogic feature, and the candidate with the highest score was selected as the annotation. Peak areas were normalised using pooled samples injected during the injection sequence. Peak area medians, standard deviations (SD), and relative SDs (RSD) were calculated using Compound Discoverer. Annotated metabolites were subsequently analysed using MetaboAnalyst 6.0^83^ using the default parameters. Significantly altered metabolites were identified for the following comparisons: ciprofloxacin versus control groups and vancomycin vs control groups. Metabolites with an FDR-corrected q-value ≤ 0.05 and an absolute fold change ≥2 were considered significantly altered.

## Acknowledgements

We thank our lab mates for helpful discussions and feedback, especially Rachel Levin and Gila Gamliel for assistance with *Drosophila* handling and experimental protocols. We further thank Kavya Nambiar and Vishakh Nair for their valuable inputs. This work was supported by the European Research Council (ERC) grant: ERC-2020-COG-BEHAVIOME-101001355.

## Data availability

The raw files for the metabolomics study were uploaded to Zenodo and will be made public upon publication (**10.5281/zenodo.20107181**). The raw files for RNA-seq were uploaded to Zenodo as well and will be made public upon publication (**10.5281/zenodo.20552513**).

## Contributions

J.D., S.T. and O.K. designed the study. J.D. performed the experiments and extractions in the study. R.S. performed the metabolomics and subsequent analysis was done by R.S. and S.K. Subsequent analyses, statistics, and visualization were done by J.D. S.T. and O.K. supervised the project. J.D. wrote the manuscript with S.T. and O.K.

## Ethics declaration

NA

